# Global Prediction of Chromatin Accessibility Using RNA-seq from Small Number of Cells

**DOI:** 10.1101/035816

**Authors:** Weiqiang Zhou, Zhicheng Ji, Hongkai Ji

## Abstract

Conventional high-throughput technologies for mapping regulatory element activities such as ChIP-seq, DNase-seq and FAIRE-seq cannot analyze samples with small number of cells. The recently developed ATAC-seq allows regulome mapping in small-cell-number samples, but its signal in single cell or samples with ≤500 cells remains discrete or noisy. Compared to these technologies, measuring transcriptome by RNA-seq in single-cell and small-cell-number samples is more mature. Here we show that one can globally predict chromatin accessibility and infer regulome using RNA-seq. Genome-wide chromatin accessibility predicted by RNA-seq from 30 cells is comparable with ATAC-seq from 500 cells. Predictions based on single-cell RNA-seq can more accurately reconstruct bulk chromatin accessibility than using single-cell ATAC-seq by pooling the same number of cells. Integrating ATAC-seq with predictions from RNA-seq increases power of both methods. Thus, transcriptome-based prediction can provide a new tool for decoding gene regulatory programs in small-cell-number samples.

## INTRODUCTION

Decoding gene regulatory network in developmental systems, precious clinical samples, and purified cells often requires measuring transcriptome (i.e., genes’ transcriptional activities) and regulome (i.e., regulatory element activities) in samples with small number of cells. While significant progress has been made to measure transcriptome in single cell (Tang et al. 2010; Ramsköld et al. 2012) and in small-cell-number (Marinov et al. 2014) samples using RNA sequencing (RNA-seq), accurately measuring regulome in single-cell and small-cell-number samples remains a challenge. Conventional high-throughput technologies such as chromatin immunoprecipitation followed by sequencing (ChIP-seq) (Johnson et al. 2007), sequencing of DNase I hypersensitive sites (DNase-seq) (Crawford et al. 2006), and Formaldehyde-Assisted Isolation of Regulatory Elements coupled with sequencing (FAIRE-seq) (Giresi et al. 2007) require large amounts of input material (˜10^6^ cells). They cannot analyze samples with small number of cells. The state-of-the-art technology ATAC-seq – assay for transposase-accessible chromatin using sequencing – can analyze chromatin accessibility in bulk samples with 500-50,000 cells (Buenrostro et al. 2013). However, ATAC-seq data are noisy when the cell number is small (e.g., ≤500). Most recently, single-cell ATAC-seq (Cusanovich et al. 2015; Buenrostro et al. 2015) (scATAC-seq) has been invented to analyze individual cells. Nevertheless, signals from scATAC-seq are intrinsically discrete since each genomic locus only has up to two copies of chromatin that can be assayed within a cell, and scATAC-seq only provides a snapshot of chromatin accessibility of a cell at the time when it is assayed and destroyed. As a surrogate for regulatory element activity, chromatin accessibility is arguably a continuous signal. This is because molecular events such as transcription-factor–DNA binding and dissociation are stochastic over time, and the overall activity of a regulatory element in a cell is determined by the probability – a continuous measure – that such stochastic events occur if one were to repeatedly observe the same cell at random time points. The discrete signal measured by scATAC-seq at a single time point cannot accurately describe this continuum of chromatin accessibility (**Supplementary Fig. 1**). Parallel to ATAC-seq, microfluidic oscillatory washing-based ChIP-seq (MOWChIP-seq) is a recently developed method for measuring histone modifications in small-cell-number samples (100-600 cells) (Cao et al. 2015). Similar to ATAC-seq, MOWChIP-seq remains noisy when the cell number is small. In a companion study, we found that chromatin accessibility measured by DNase I hypersensitivity (DH) in a bulk sample can be predicted with good accuracy using the sample’s gene expression profile measured by Affymetrix exon array (Zhou et al. submitted). We also developed a computational method BIRD to handle this big data prediction problem (**Supplementary Methods, Supplementary Fig. 2**). Here we investigate whether one can use a similar approach to predict regulome based on RNA-seq, and importantly, whether this approach allows one to use small-cell-number and single-cell RNA-seq (scRNA-seq) to predict regulome in samples with limited amounts of materials (**Fig. 1a**).

**Figure 1.**
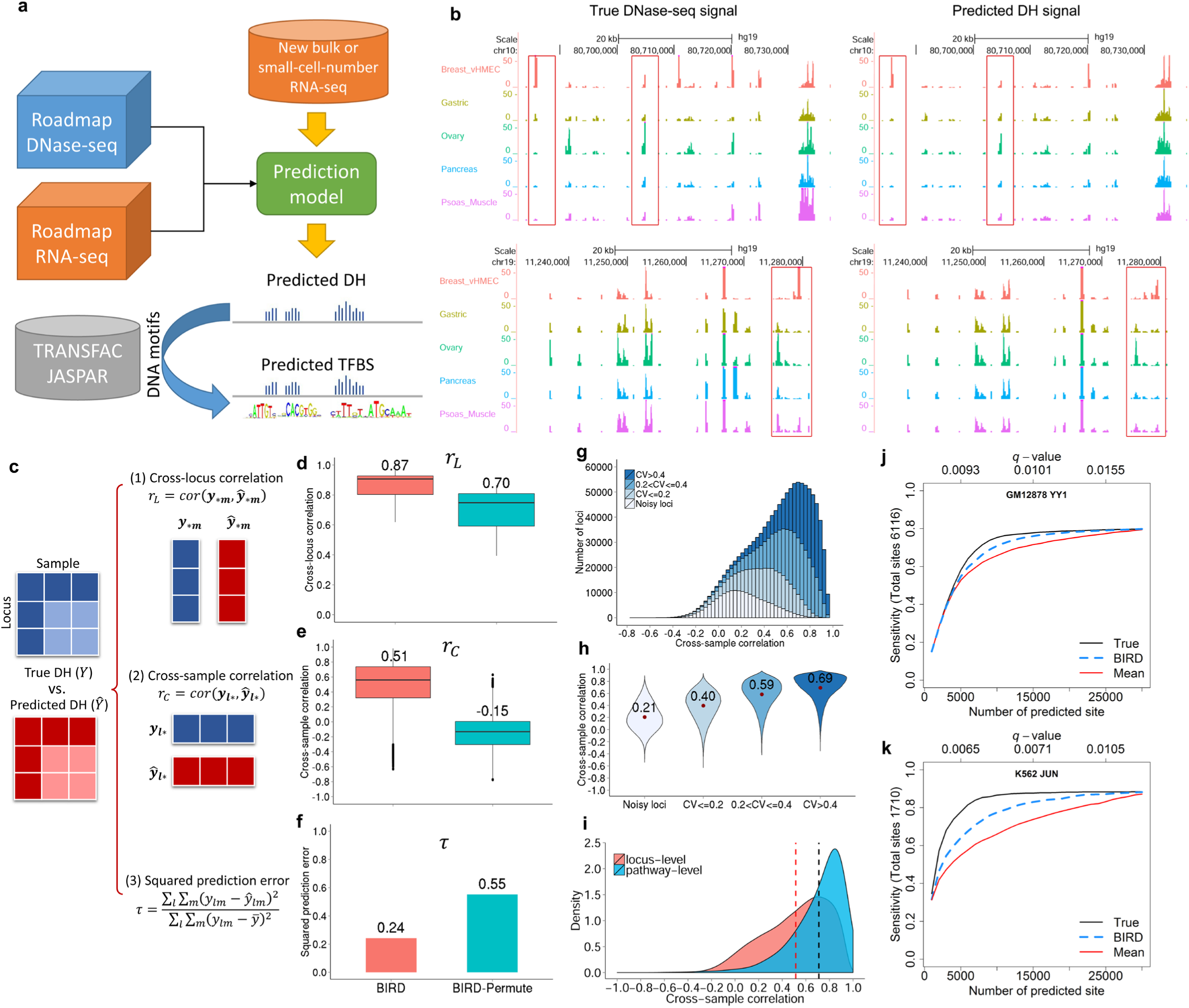
BIRD predicts DH and TFBSs using bulk RNA-seq. (**a**) Overview of the study. Roadmap Epigenomics DNase-seq and RNA-seq data are used to train BIRD prediction models which are then applied to new RNA-seq samples to predict DH. The predicted DH can be coupled with DNA motifs to predict TFBSs. (**b**) Two examples of true and predicted DH signals across five different samples. Each track is a sample. Regions highlighted with boxes demonstrate that the predicted DH captures the true DH variation. (**c**) Statistics used to evaluate prediction performance. (**d**)-(**f**) Prediction performance of BIRD and random prediction models (“BIRD-permute”) in leave-one-cell-type-out cross-validation. (**d**) Distribution and mean of cross-locus correlation *r_L_* from all samples. (**e**) Distribution and mean of cross-sample correlation *r_C_* from all loci. (**f**) Squared prediction error (*τ*). (**g**) Genomic loci are grouped into four categories by coefficient of variation (CV) of the predicted DH across samples at each locus. Distribution of *r_C_* of all loci, stratified using the four CV categories, is shown for BIRD. (**h**) Distribution and mean of *r_C_* in each CV category. (**i**) Distribution of *r_C_* for locus-level predictions vs. pathway-level predictions. (**j**)-(**k**) Sensitivity-rank curve for predicting YY1 binding sites in GM12878 and JUN binding sites in K562 cells using true DNase-seq (“True”), BIRD, and mean DH profile of training samples (“Mean”). For each method, the curve shows the percentage of true TFBSs discovered by top predicted motif sites. *q*-values corresponding to top 5000, 15000, and 25000 BIRD predictions are shown on top of each plot.

## RESULTS

### Predicting Chromatin Accessibility Using Bulk RNA-seq

We begin with evaluating the feasibility of using bulk RNA-seq to predict DH. We downloaded DNase-seq and matching RNA-seq data for 70 human samples representing 30 different cell types (**Supplementary Table 1**) from the Roadmap Epigenomics project (Kundaje et al. 2015) (also called Epigenome Roadmap below). After preprocessing and normalization, 37,335 transcripts with expression measurements from RNA-seq and 1,136,465 genomic loci with DH measurements from DNase-seq were obtained and served as predictors and responses respectively (**Methods**). Our goal is to predict DH at these 1,136,465 loci using the 37,335 predictors. We evaluate the prediction using leave-one-out cross-validation. In each fold of the cross-validation, the 30 cell types were partitioned into a training dataset consisting of 29 cell types and a test dataset consisting of 1 cell type. BIRD prediction models were trained using samples in the training data and then applied to RNA-seq samples in the test data to predict DH (**Methods**). Prediction performance was evaluated using true DNase-seq signals in the test data and the following statistics (**Fig. 1c**): (1) Pearson correlation between the predicted and true DH values across all genomic loci within each sample (“cross-locus correlation” *r_L_*), (2) Pearson correlation between the predicted and true DH values across all samples at each genomic locus (“cross-sample correlation” *r_C_*), and (3) total squared prediction error scaled by the total DH data variance (r). As a control, we also constructed random prediction models (“BIRD-Permute”) by permuting the link between the DNase-seq and RNA-seq samples in the training data and then applied them to the test data.

Based on *r_L_*, RNA-seq was able to accurately predict how DH varied across different genomic loci (mean *r_L_*=0.87), and the prediction accuracy of BIRD was significantly higher than random expectation (**Fig. 1d**, BIRD vs. BIRD-Permute: two-sided Wilcoxon signed-rank test *p*-value = 3.6×10^−13^). Of note, BIRD-Permute also explained a large amount of cross-locus DH variation (**Fig. 1d**, mean *r_L_* = 0.70). This was caused by strong locus-dependent DH propensities not perturbed by permutation (**Supplementary Fig. 3, Methods**). Due to these locus effects, simply using the mean DH profile across training samples can predict cross-locus DH variation to certain extent (**Supplementary Fig. 4**), although such prediction is independent of test sample and therefore less accurate compared to BIRD predictions which utilize test-sample-dependent transcriptome information.

Based on *r_C_*, RNA-seq was also able to predict how DH varied across samples with substantially higher accuracy than random expectation (**Fig. 1e**, mean *r_C_* of BIRD vs. BIRD-Permute = 0.51 vs. −0.15, two-sided Wilcoxon signed-rank test *p*-value < 2.2×10^−16^). **Figure 1b** shows two examples of such prediction. Prediction of cross-sample variation was less accurate than prediction of cross-locus variation (**Fig. 1d-e**, BIRD mean *r_C_* vs. mean *r_L_* = 0.51 vs. 0.87), because cross-sample prediction performance was evaluated within each locus and not affected by the locus effects. Cross-sample prediction accuracy varied substantially across loci (**Fig. 1e**). A large proportion of loci can be predicted with good accuracy: 57% and 23% of loci had *r_C_* > 0.5 and >0.75, respectively. For each locus, the coefficient of variation (CV) of the predicted DH values across samples was computed to characterize its cross-sample DH variability (Methods). It was observed that loci with smaller *r_C_* also tend to have smaller CV (**Fig. 1g**). On average, prediction of cross-sample variability was more accurate for loci with higher variability (**Fig. 1h**). For instance, for loci with CV>0.4, the mean *r_C_* was 0.69 (>0.51, the mean *r_C_* of all loci), and 84% and 47% of such loci had *r_C_* > 0.5 and >0.75 respectively.

By grouping genomic loci with similar cross-sample DH variation patterns into clusters and treating each cluster as a “pathway” of co-activated regulatory elements (**Methods**), cross-sample variation of the pathway activity (i.e., mean DH of all loci in each pathway) can be predicted more accurately (mean *r_C_*=0.71, 85% and 55% of pathways had *r_C_* > 0.5 and >0.75) than predicting cross-sample variability of individual loci (**Fig. 1i**).

BIRD prediction also substantially reduced the squared prediction error compared to random expectation (**Fig. 1f**, BIRD vs. BIRD-Permute; *τ*=0.24 vs. 0.55). Together, the above results are consistent with the results in the companion study where DH was predicted using exon arrays (Zhou et al. submitted).

### Predicting Transcription Factor Binding Sites Using Bulk RNA-seq

We tested if the predicted DH at DNA motif sites can predict transcription factor (TF) binding sites (TFBSs) by analyzing 34 TFs in GM12878 and 25 TFs in K562 cells. BIRD models trained using the Epigenome Roadmap data (70 samples, GM12878 and K562 were not part of the 70 samples) were applied to predict DH in GM12878 and K562 using RNA-seq. The DNA motif of each TF was mapped to the genome, and motif sites with high predicted DH were identified and ranked as predicted TFBSs. For each TF and cell type, the corresponding ChIP-seq data were obtained from ENCODE (ENCODE Project Consortium 2012). Motif-containing ChIP-seq peaks were used as gold standard to evaluate the prediction accuracy (**Methods**). **Figure 1j-k** and **Supplementary Figures 5-6** show the percentage of gold standard TFBSs that were discovered by the top predicted sites. For comparison, we also predicted TFBSs using the true DNase-seq data (positive control) and the mean DH profile of the training samples (negative control). The results show that BIRD predictions based on RNA-seq were able to discover a substantial proportion of the true TFBSs. For instance, the top 15,000 predictions for YY1 in GM12878 (*q*-value = 0.01) covered 76% of the gold standard YY1 binding sites (**Fig. 1j**). As expected, predictions based on true DNase-seq were more accurate than BIRD predictions. However, compared to the cell-type-independent prediction based on the mean DH profile, BIRD predictions were substantially better because BIRD used cell-type-specific information contained in the transcriptome.

### Predicting Chromatin Accessibility Using Small-cell-number RNA-seq

Our next question is whether BIRD trained using bulk RNA-seq data from the Epigenome Roadmap can be applied to RNA-seq generated using small-cell-number samples to predict DH. We obtained published bulk RNA-seq and RNA-seq from samples with 10, 30 and 100 cells for GM12878 (Marinov et al. 2014). BIRD models trained using the Epigenome Roadmap data were applied to each sample. For evaluation, true chromatin accessibility profiles for these small-cell-number samples are not available. However, according to statistical theory, if cells in a small-cell-number sample are randomly drawn from a bulk cell population, the mean DH profile of the small-cell-number sample and that of the bulk sample should have the same expectation. Therefore, one can use the bulk DNase-seq data for GM12878 (ENCODE Project Consortium 2012) from the ENCODE as the “truth”. Based on this gold standard, we compared BIRD predictions with GM12878 ATAC-seq from 500 and 50,000 cells. Our evaluation was primarily based on cross-locus correlation *r_L_*, because reliably estimating cross-sample correlation *r_C_* requires a large number of test cell types which were not available here. It turns out that ATAC-seq from 50,000 cells (“ATAC-b50k”) showed the highest cross-locus correlation with the true DNase-seq signal (**Fig. 2a-b**, *r_L_*=0.76). Surprisingly, however, BIRD-predicted DH signals from 30 and 100 cells consistently predicted the truth better than ATAC-seq from 500 cells (**Fig. 2a-b**, *r_L_*=0.63, 0.69 and 0.69 for “ATAC-b500”, “BIRD-b30” and “BIRD-b100”). Of note, using the mean DH profile from the training data alone was able to predict DH to certain degree (**Fig. 2a-b**, *r_L_*=0.56 for “Mean”). The prediction accuracy of BIRD increased with increasing cell number. BIRD predictions based on ≥30 cells were almost as accurate as predictions based on bulk RNA-seq (**Fig. 2a-b**, *r_L_*=0.70 for “BIRD-bulk”). **Figure 2b** provides an example illustrating signals from different methods.

**Figure 2.**
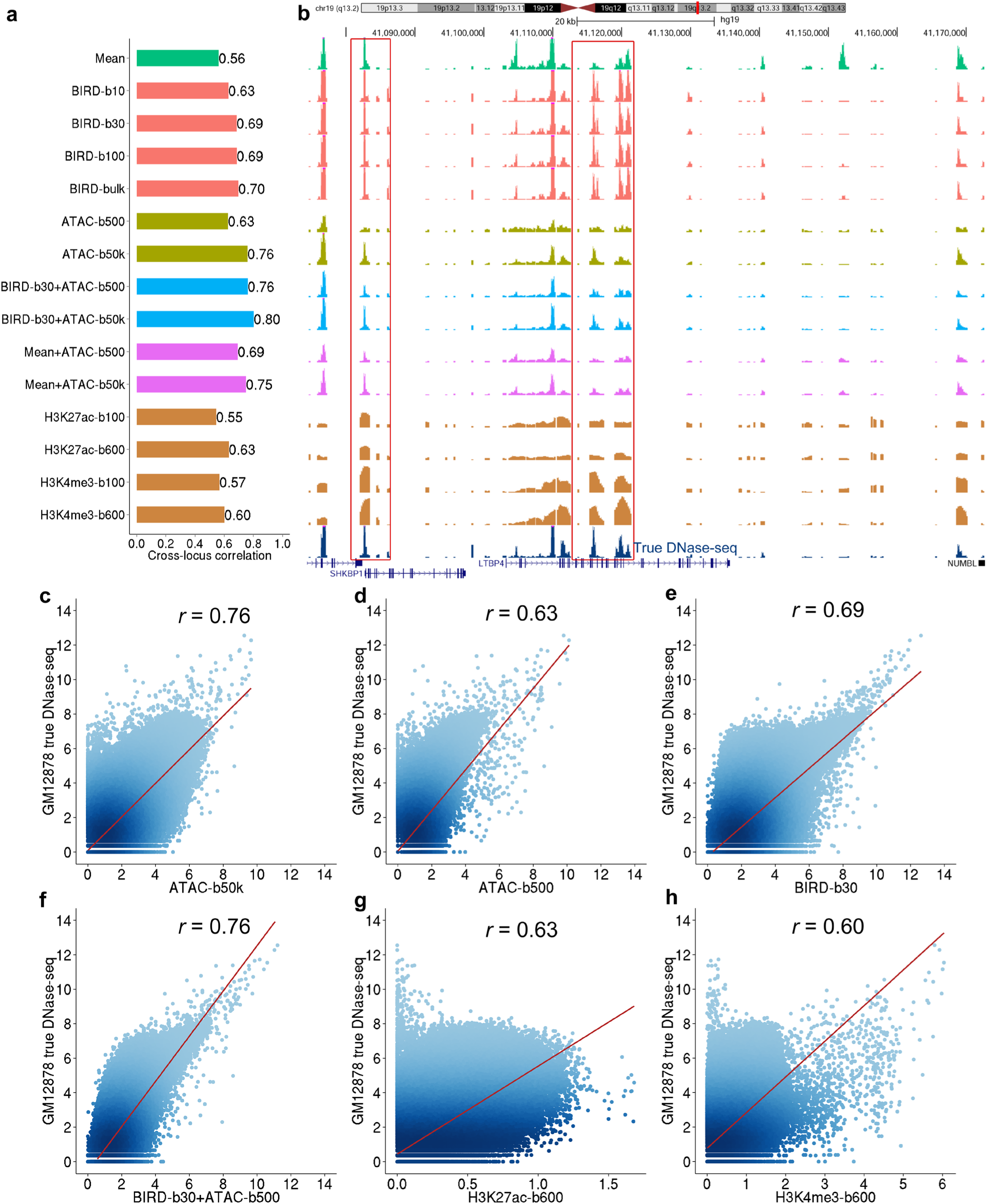
Predicting DH using small-cell-number RNA-seq data. (**a**) Cross-locus correlation between the bulk GM12878 DNase-seq signal and chromatin accessibility predicted or measured by different methods. “Mean”: mean DH profile of training samples. “BIRD-b10”, “BIRD-b30”, “BIRD-b100”: BIRD-predicted DH based on small-cell-number RNA-seq samples with 10, 30 and 100 cells. “BIRD-bulk”: BIRD-predicted DH based on bulk RNA-seq. “ATAC-b500”, “ATAC-b50k”: ATAC-seq with 500 and 50,000 cells. “BIRD-b30+ATAC-b500”, “BIRD-b30+ATAC-b50k”: average of BIRD-predicted DH from 30 cells and ATAC-seq from 500 or 50,000 cells. “Mean+ATAC-b500”, “Mean+ATAC-b50k”: average of mean DH profile of training samples and ATAC-seq from 500 or 50,000 cells. “H3K27ac-b100”, “H3K27ac-b600”, “H3K4me3-b100” and “H3K4me3-b600”: MOWChIP-seq for histone modification H3K27ac or H3K4me3 with 100 or 600 cells. (**b**) An example that compares chromatin accessibility predicted or measured by different methods. True bulk DNase-seq signal is shown on the bottom track as a reference. Regions highlighted by boxes illustrate that BIRD predicted DH better than “Mean” and “ATAC-b500”. (**c**)-(**h**) Scatterplots comparing true bulk DNase-seq signal with chromatin accessibility predicted or measured by ATAC-b50k, ATAC-b500, BIRD-b30, BIRD-b30+ATAC-b500, H3K27ac-b600 and H3K4me3-b600. Each dot is a genomic locus. The cross-locus correlation is shown on top of each plot.

Interestingly, combining the ATAC-seq signal from 500 cells and the BIRD-predicted DH from 30 cells by average (530 cells used in total) allowed one to better predict the gold standard DNase-seq signal (**Fig. 2a-b**, “BIRD-b30+ATAC-b500”, *r_L_*=0.76). The combined signal achieved the same accuracy as ATAC-seq using 50,000 cells (*r_L_*=0.76) and was better than using BIRD-b30 (*r_L_*=0.69) or ATAC-b500 (*r_L_*=0.63) alone (**Fig. 2c-f**). Similarly, by averaging ATAC-seq from 50,000 cells and BIRD predictions from 30 cells, we were able to predict the gold standard better than ATAC-b50k (**Fig. 2a-b**, *r_L_*=0.80 for “BIRD-b30+ATAC-b50k”). The same improvement was not observed when the BIRD prediction was replaced by the prediction based on the mean DH profile (**Fig. 2a-b**, *r_L_*=0.69 and 0.75 for “Mean+ATAC-b500” and “Mean+ATAC-b50k”). These results show that DH predicted from small-cell-number RNA-seq can be integrated with small-cell-number ATAC-seq data (BIRD+ATAC-seq) to obtain better signal.

We repeated the above evaluation by using ATAC-seq from 50,000 cells to replace bulk DNase-seq to serve as gold standard. Similar conclusions were obtained (**Methods, Supplementary Fig. 7**). Unlike the DNase-seq gold standard which came from a study different from the studies that generated the test ATAC-seq and RNA-seq data, the ATAC-50k gold standard was collected from the same study as ATAC-b500 (RNA-seq was from a different study). Thus, the ATAC-50k gold standard should intrinsically favor ATAC-b500 over BIRD due to potential lab effects. Despite this, BIRD predictions based on ≥30 cells performed close to ATAC-b500 in this comparison, and BIRD-b30+ATAC-b500 outperformed ATAC-b500 (**Supplementary Fig. 7**).

### Predicting Transcription Factor Binding Sites Using Small-cell-number RNA-seq

We further evaluated whether DH predicted using small-cell-number RNA-seq coupled with DNA motif information can predict TFBSs. Similar to the analyses performed for the bulk RNA-seq, we predicted TFBSs for 34 TFs in GM12878 using BIRD-b30, BIRD-hybrid (i.e. BIRD-b30+ATAC-b500), ATAC-b50k, ATAC-b500, true DNase-seq (“True”), and mean DH of training samples (“Mean”). **Figure 3a-f** and **Supplementary Figure 8** show the performance curves. To facilitate method comparison, we calculated the area under the curve (AUC) for each method, normalized by dividing the AUC of the “True” DNase-seq (**Fig. 3g**, **Supplementary Table 3, Methods**). Comparison of the normalized AUC shows that BIRD-b30 outperformed mean DH in all 34 tested TFs. Furthermore, BIRD-b30 outperformed ATAC-b500 in 23 of 34 TFs (**Fig. 3g**, **Supplementary Fig. 8**). Interestingly, BIRD-hybrid (BIRD-b30+ATAC-b500) outperformed ATAC-b500 in 32 of 34 TFs, and outperformed ATAC-b50k in 21 of 34 TFs. Thus, DH predicted by BIRD from 30 cells more accurately predicted TFBSs than ATAC-seq from 500 cells, and combining BIRD predictions with ATAC-seq from small number of cells better predicted TFBSs than bulk ATAC-seq.

**Figure 3.**
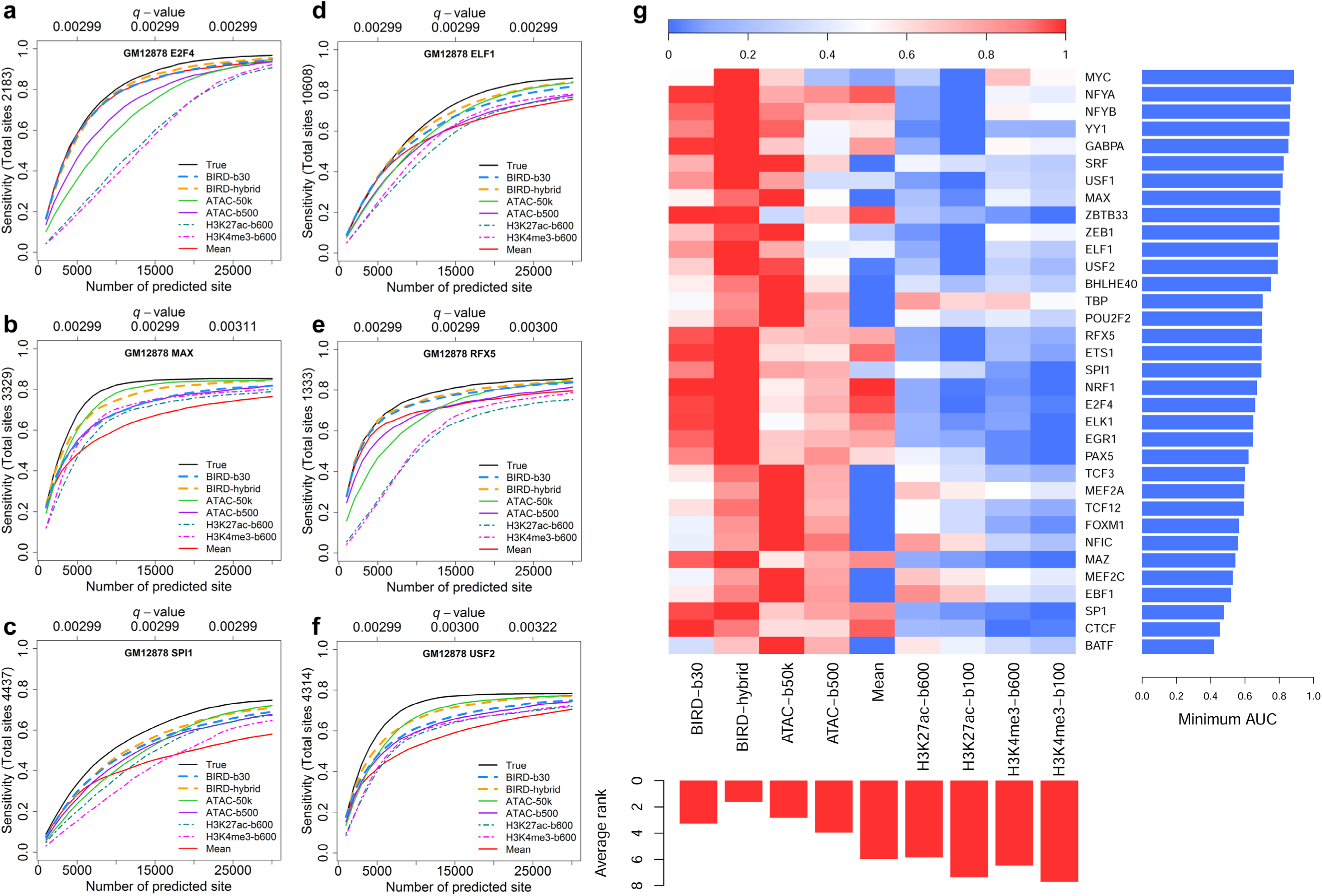
Predicting TFBSs using small-cell-number RNA-seq data. (**a**)-(**f**) Sensitivity-rank curve for predicting E2F4, MAX, SPI1, ELF1, RFXS and USF2 binding sites in GM12878 using true DNase-seq (“True”), ATAC-seq from 500 or 50,000 cells (“ATAC-500”, “ATAC-b50k”), mean DH profile of training samples (“Mean”), BIRD-predicted DH using 30 cells (“BIRD-b30”), the average of BIRD-predicted DH using 30 cells and ATAC-seq using 500 cells (“BIRD-hybrid”), and MOWChIP-seq for H3K27ac and H3K4me3 using 600 cells (“H3K27ac-b600”, “H3K4me3-b600”). The performance for MOWChIP-seq using 100 cells was generally worse than using 600 cells and hence is shown in **Supplementary Figure 8** but not shown here for clarity of display. The *q*-values for BIRD-b30 predictions are shown on the top of each plot. (**g**) Scaled area under the curve (AUC) for different methods in TFBS prediction. Each row is a TF, and each column is a method. For each TF, different methods are ranked based on the AUC value, and the worst AUC value of all methods is shown on the right using a blue bar. The average rank of each method across all TFs is shown on the bottom using a red bar. Smaller rank means better performance.

### A Comparison of BIRD, ATAC-seq and MOWChIP-seq for Small-cell-number Samples

Next, we compared DH predicted by BIRD using 30 cells, ATAC-seq, and histone modification H3K27ac and H3K4me3 profiles measured by MOWChIP-seq using 100 and 600 GM12878 cells. Since the genomic distribution of histone modification signal is different from that of chromatin accessibility due to nucleosome displacement around TFBSs (He et al. 2010), we first optimized the parameter for analyzing MOWChIP-seq data (**Methods, Supplementary Fig. 9**). The comparisons below are based on the optimal MOWChIP-seq performance. It was observed that predictions or measurements for each data type correlated better with the bulk data from the same data type than the bulk data from other data types (**Supplementary Fig. 10**). For instance, H3K27ac MOWChIP-seq using 100 and 600 cells (H3K27ac-b100 and H3K27ac-b600) performed better than BIRD-b30 when H3K27ac bulk ChIP-seq was used as gold standard for evaluation, but the same MOWChIP-seq data performed worse than BIRD-30 when bulk DNase-seq was used as gold standard (**Supplementary Fig. 10, Fig. 2a,b,g,h**). This suggests that there were substantial differences among data types, making a fair comparison difficult. For predicting TFBSs, however, both BIRD-b30 and ATAC-b500 substantially outperformed MOWChIP-seq based on the overall performance in all 34 tested TFs (**Fig. 3a-f**, **Supplementary Fig. 8**). Among the MOWChIP-seq data, H3K27ac-b600 had the best overall performance for predicting TFBSs (**Fig. 3g**, **Supplementary Table 3**). BIRD-b30 outperformed H3K27ac-b600 MOWChIP-seq in 27 of 34 tested TFs. ATAC-b500 outperformed H3K27ac-b600 in 31 of 34 tested TFs. Finally, BIRD-hybrid (BIRD-b30+ATAC-b500) outperformed H3K27ac-b600 in 33 out of 34 TFs.

**Figure 4.**
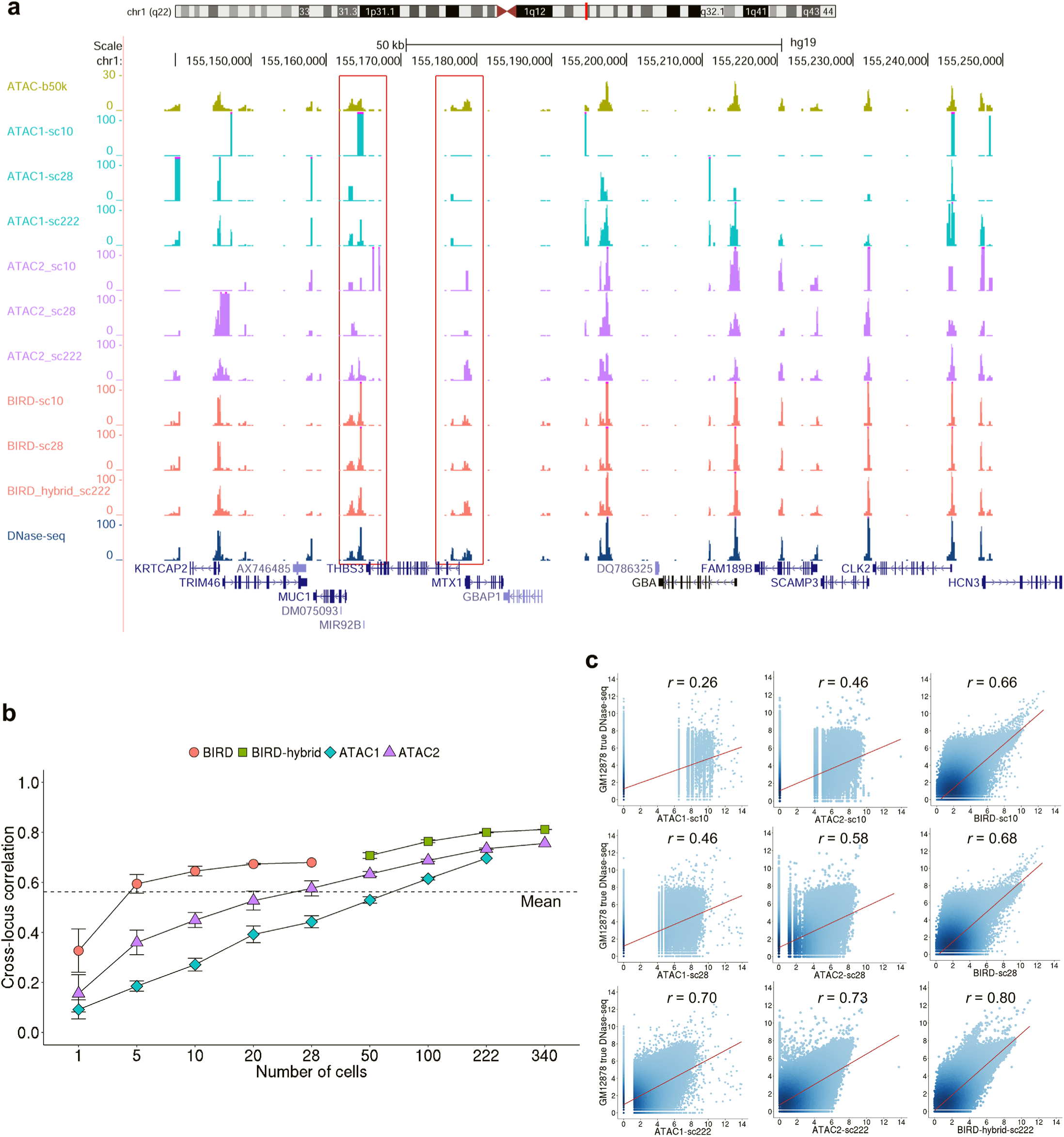
BIRD predicts DH using pooled single-cell RNA-seq data. (**a**) An example comparing chromatin accessibility reported by different single-cell methods. “ATAC1-sc10”, “ATAC1-sc28” and “ATAC1-sc222”: pooled single-cell ATAC-seq from 10, 28 or 222 cells using scATAC-seq dataset 1. “ATAC2-sc10”, “ATAC2-sc28” and “ATAC2-sc222”: pooled single-cell ATAC-seq from 10, 28 or 222 cells using scATAC-seq dataset 2. “BIRD-sc10”, “BIRD-sc28”: BIRD-predicted DH based on pooled single-cell RNA-seq data from 10 or 28 cells. “BIRD-hybrid-sc222”: the average of BIRD-sc28 and single-cell ATAC-seq from 194 cells using scATAC-seq dataset 2. As references, bulk ATAC-seq from 50,000 cells (“ATAC-b50k”) and DNase-seq are shown on the top and bottom respectively. (**b**) Cross-locus correlation between the true bulk DNase-seq signal and chromatin accessibility predicted or measured by different single-cell methods. The correlation is shown as a function of pooled cell number. Error bars are standard deviation based on 10 independent samplings of cells (Methods). “ATAC1”: scATAC-seq dataset 1. “ATAC2”: scATAC-seq dataset 2. “BIRD”: BIRD-predicted DH using pooled single-cell RNA-seq. “BIRD-hybrid”: the average of BIRD-predictions based on 28 cells and pooled ATAC-seq from scATAC-seq dataset 2 (here x-axis is the total number of cells used by scRNA-seq and scATAC-seq). Prediction performance using the mean DH profile of training samples (“Mean”) is shown as a dashed line. (**c**) Scatterplots comparing true bulk DNase-seq signal with chromatin accessibility predicted or measured by ATAC1, ATAC2 and BIRD (or BIRD-hybrid for 222 cells) using 10, 28 and 222 cells. Each dot is a genomic locus. The cross-locus correlation is shown on top of each plot.

### Predicting Chromatin Accessibility and TFBSs Using Single-cell RNA-seq

We proceeded to investigate whether one can use single-cell RNA-seq data to predict DH. We analyzed a single-cell RNA-seq dataset with 28 single cells for GM12878 (Marinov et al. 2014). After calculating gene expression for each cell, we pooled *k* (*k* = 1, 5, 10, 20, 28) cells randomly drawn from the dataset together and used their average expression profile to predict DH based on BIRD models trained from the Epigenome Roadmap bulk RNA-seq data. For comparison, we analyzed published single-cell ATAC-seq data in GM12878 generated by two different protocols (“ATAC1” (Cusanovich et al. 2015): 222 cells; “ATAC2” (Buenrostro et al. 2015): 340 cells). We computed average scATAC-seq profile for *k* (*k* = 1, 5, 10, 20, 28, 50, 100, 222 and 340) cells randomly drawn from each dataset respectively. **Figure 4** shows the performance of different methods evaluated using bulk DNase-seq as gold standard. Holding the cell number the same, BIRD based on pooled scRNA-seq was consistently better than pooled scATAC-seq for predicting bulk DNase-seq (**Fig. 4b,c**). BIRD predictions based on a single cell and pooled scATAC-seq using ≤50 cells from ATAC1 or ≤20 cells from ATAC2 were less accurate than predictions based on the mean DH profile (**Fig. 4b**). However, prediction accuracy increased as more cells were pooled together. BIRD with 10 cells performed better than the mean DH profile, and it was comparable to pooling 100 cells from ATAC1 or pooling 50 cells from ATAC2. The results remained similar when the gold standard was changed to bulk ATAC-seq data from 50,000 or 500 cells (**Supplementary Fig. 11**).

We also combined BIRD predictions based on pooling scRNA-seq from 28 cells with the pooled scATAC-seq profile from *x* cells (*x* = 22, 72, 194 and 312) by taking the average of the two profiles (“BIRD-hybrid”). We then compared BIRD-hybrid with pooled scATAC-seq data using the same number of cells (i.e., *k* = 28 + *x* = 50, 100, 222, 340). BIRD-hybrid also outperformed pooled scATAC-seq (**Fig. 4a-c, Supplementary Fig. 11**).

To test whether predictions from scRNA-seq can predict TFBSs in a similar fashion as small-cell-number RNA-seq, we again analyzed 34 TFs in GM12878 (**Fig. 5a-f, Supplementary Figs. 12-15**, **Supplementary Table 4**). Once again, BIRD and BIRD-hybrid performed better than pooled scATAC-seq. For instance, when pooling 10 cells, BIRD prediction outperformed ATAC1 and ATAC2 in all 34 TFs, and it outperformed mean DH in 33 of 34 TFs. Using 28 cells, BIRD outperformed ATAC1, ATAC2 and mean DH in 34, 32 and 33 out of 34 tested TFs respectively. Using 222 single cells, BIRD-hybrid outperformed ATAC1, ATAC2 and mean DH in 33, 33 and 31 out of the 34 tested TFs respectively (**Supplementary Fig. 15**, **Supplementary Table 4**).

**Figure 5.**
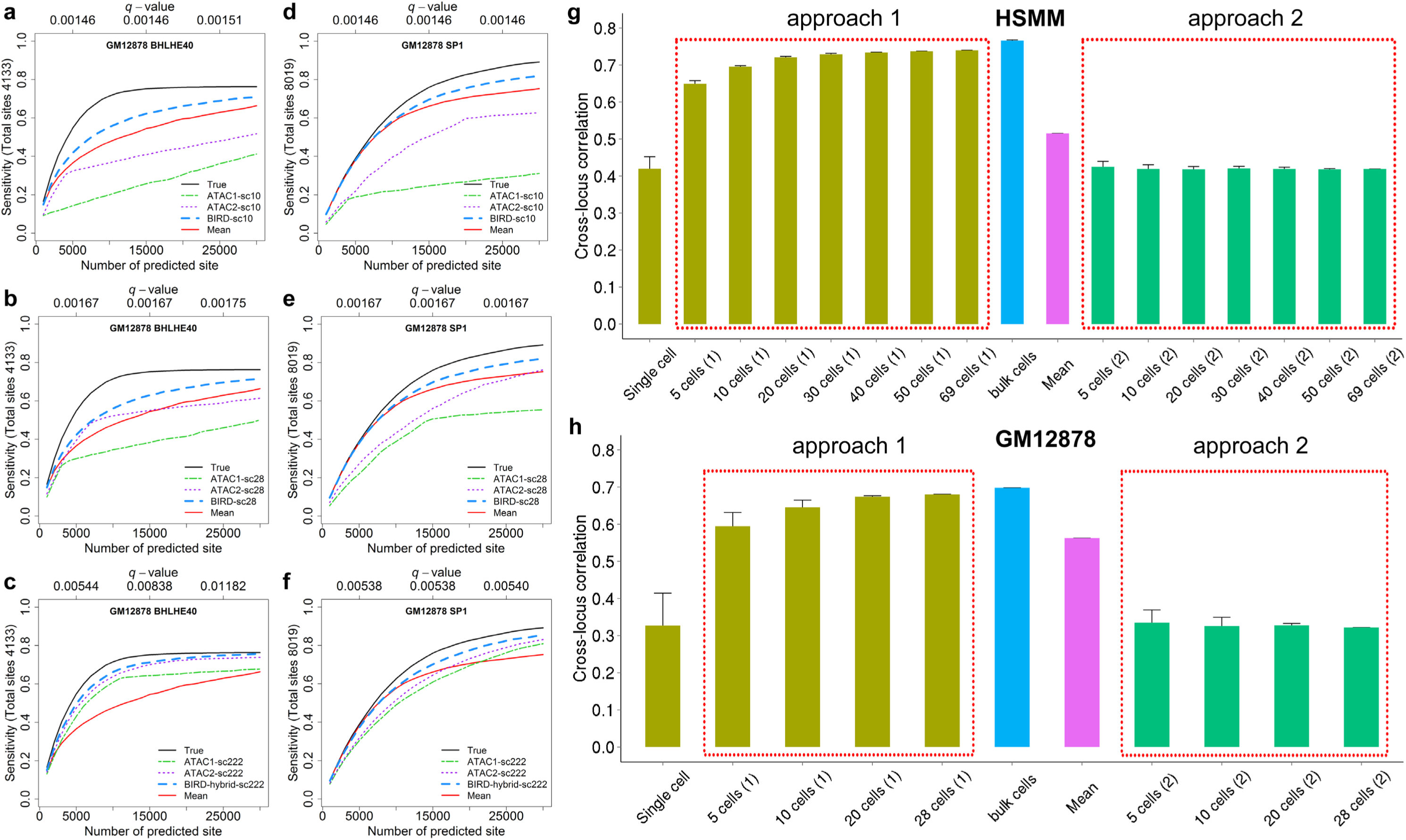
BIRD predicts TFBSs using pooled single-cell RNA-seq data and a comparison between two different prediction strategies. (**a**)-(**f**) Sensitivity-rank curve for predicting BHLHE40 and SP1 binding sites in GM12878 using true DNase-seq (“True”), mean DH profile of training samples (“Mean”), and BIRD and scATAC-seq by pooling different number of cells. (**a,d**) Pooled scATAC-seq and BIRD using 10 cells. (**b,e**) Pooled scATAC-seq and BIRD using 28 cells. (**c,f**) Pooled scATAC-seq and BIRD-hybrid using a total of 222 cells. “ATAC1” and “ATAC2” correspond to two different scATAC-seq datasets. *q*-values shown in each plot are calculated based on BIRD-sc10, BIRD-sc28 and BIRD-hybrid-sc222 predictions, respectively. (**g**) Cross-locus correlation between the true bulk DNase-seq signal and BIRD-predicted DH using two different prediction strategies in the HSMM dataset. Approach 1 (“5 cells(1)”,…, “69 cells(1)”): pool scRNA-seq from multiple cells first and then use the pooled scRNA-seq to make predictions; Approach 2 (“5 cells(2)”,…, “69 cells(2)”): use scRNA-seq from each single cell to make prediction first and then pool predictions from different cells by averaging. Error bars are standard deviation based on 10 independent samplings of cells (Methods). (**h**) Cross-locus correlation between the true bulk DNase-seq signal and BIRD-predicted DH using two different prediction strategies in GM12878. Approach 1 (“5 cells(1)”,…, “28 cells(1)”) and Approach 2 (“5 cells(2)”,…, “28 cells(2)”) are the same as above.

We applied BIRD to another scRNA-seq dataset (Trapnell et al. 2014) (69 cells) from human skeletal muscle myoblasts (HSMM) (**Fig. 5g**, “approach 1”, pooling *k* = 1, 5, 10, 20, 30, 40, 50 and 69 cells). For this dataset, scATAC-seq was not available and therefore not compared. We used bulk DNase-seq data in HSMM as gold standard for evaluation. The prediction accuracy of BIRD by pooling ≥5 cells was better than the accuracy based on the mean DH profile, and the accuracy of BIRD by pooling ≥30 cells was comparable to BIRD predictions based on bulk RNA-seq (**Fig. 5g**). This further demonstrates that one can predict DH from scRNA-seq by pooling a small number of cells.

When applying BIRD to scRNA-seq data, it is important to pool RNA-seq data from multiple cells first and then make predictions based on the pooled gene expression profile. When we tried to first predict DH based on each single cell and then average the predictions from multiple cells, the prediction performance was substantially worse for both the GM12878 and HSMM data (**Fig. 5g-h**, “approach 2”). This is because expression measurements from scRNA-seq have substantial biases (e.g., zero-inflation by dropout events (Kharchenko et al. 2014)) that cannot be removed by the usual normalization. Impacts on prediction by such bias can be reduced when multiple cells are pooled together to measure gene expression and the pooled expression profile is then normalized against the bulk RNA-seq data in the training dataset.

## DISCUSSION

To summarize, our analyses demonstrate that predicting chromatin accessibility using RNA-seq can provide a new approach for regulome mapping both in bulk samples and in samples with small number of cells. The study compared multiple state-of-the-art technologies for mapping regulome in small-cell-number samples including ATAC-seq, scATAC-seq, MOWChIP-seq and BIRD. Our results show that for analyzing small-cell-number samples, BIRD can offer competitive performance compared to ATAC-seq and scATAC-seq. In particular, using 5-10 folds fewer cells, BIRD reached the same accuracy as ATAC-seq and pooled scATAC-seq for predicting bulk chromatin accessibility. Also, BIRD based on scRNA-seq more accurately predicted bulk chromatin accessibility than using scATAC-seq by pooling the same number of cells. Besides ATAC-seq, BIRD based on fewer cells also offered competitive or better performance compared to MOWChIP-seq using more cells.

Based on our analyses, the minimum number of cells required by the current technology to recover chromatin accessibility in a bulk sample is approximately 10 cells. This was achieved by BIRD. Averaging single-cell ATAC-seq from 10 cells predicted bulk chromatin accessibility worse than the trivial prediction based on the mean DH profile. By contrast, BIRD predictions based on pooling scRNA-seq from 10 cells were better than predictions based on the mean DH profile. This highlights the limitation of scATAC-seq due to its intrinsic discreteness (**Supplementary Fig. 1**). Compared to scATAC-seq, scRNA-seq data are less discrete since each gene can have more than two copies of transcripts in a cell.

Most recently, single-cell ChIP-seq (Drop-ChIP) for histone modifications and single-cell DNase-seq (scDNase-seq) have been reported (Rotem et al. 2015; Jin et al. 2015). Since the current Drop-ChIP and scDNase-seq data for single cells are in mouse and we do not have enough training samples in mouse to build BIRD models, we were unable to directly compare BIRD with Drop-ChIP and scDNase-seq here. Of note, Drop-ChIP data are highly discrete, with 500˜10,000 reads and an average of ˜800 peaks detected per cell. Our results on scATAC-seq (ATAC1 and ATAC2 had an average of ˜2,700 and ˜14,000 reads per cell respectively) suggest that discreteness of the signal will remain a problem for Drop-ChIP. Although scDNase-seq has also been applied to pooled human cells dissected from formalin-fixed paraffin-embedded tissues, there is no gold standard available for a direct comparison between BIRD and scDNase-seq for that application. In the future, it will be interesting to compare BIRD with Drop-ChIP and scDNase-seq when appropriate test and benchmark data become available.

Our study has important practical relevance on future data analyses. It shows that transcriptome-based regulome prediction can greatly increase the value of current and future bulk, small-cell-number and single-cell RNA-seq experiments. By adding a new component to the standard RNA-seq analysis pipeline, this approach allows one to use RNA-seq not only for studying transcriptome but also for studying regulome. This can greatly impact how to most effectively use existing and future RNA-seq data, which is particularly relevant given that enormous amounts of RNA-seq data will be generated in the years to come.

Our study also has important implications for future experiment design. When a sample contains only a very limited number of cells, researchers have to decide how these cells should be wisely used. For example, should one use all cells for transcriptome profiling by RNA-seq or regulome mapping by ATAC-seq? Results from this study show that one may divide the samples into two parts, one for RNA-seq or scRNA-seq, and one for ATAC-seq or scATAC-seq. This strategy has two advantages. First, one can obtain information for two different data types instead of only one data type. Second, by spending some cells on RNA-seq, BIRD-hybrid allows one to combine the two data types to produce comparable or better regulome mapping than spending all cells on ATAC-seq. This study also shows that if one decides to use all cells for RNA-seq, one can still obtain information on regulome through prediction. Thus, it is also possible to analyze transcriptome and regulome simultaneously in a small-cell-number sample by measuring only transcriptome.

Currently, BIRD predictions based on RNA-seq from a single cell were less accurate than the mean DH profile for predicting the bulk chromatin accessibility. One possible reason is that technical biases in the single-cell RNA-seq data (e.g., excessive zeros in the data) cannot be easily removed by normalization when there is only one cell, making the prediction inaccurate. Another possible reason is that the small sample size (n=1 cell) is not sufficient to overcome the random variation in single-cell expression to recover the behavior of a bulk sample. Naturally, an important question for future research is whether one can develop methods insensitive to the biases in scRNA-seq to improve predictions in a single cell. More generally, there is still great demand for new experimental or computational methods for single-cell regulome mapping, particularly in the context that the discrete signals generated at one random time point by current experimental technologies such as scATAC-seq may not adequately describe the average steady-state behavior of a cell over time.

As a proof-of-concept, this study shows that predicting chromatin accessibility using bulk and small-cell-number RNA-seq is feasible. Another important next step is to explore whether other functional genomic data types can be predicted in a similar fashion in small-cell-number samples.

## METHODS

### DNase-seq data processing

The aligned DNase-seq data (alignment based on hg19) from 70 samples were downloaded from the Roadmap Epigenomics project (Kundaje et al. 2015) (ftp://ftp.genboree.org/EpigenomeAtlas/Current-Release/experiment-sample/Chromatin_Accessibility/). The analyses in this study were focused on chromosomes 1 to X. Excluding chromosome Y, the genome was divided into 200 base pair (bp) non-overlapping bins. The number of reads mapped to each bin was counted for each DNase-seq sample. To adjust for different sequencing depths, bin read counts for each sample *i* were first divided by the sample’s total read count *N_i_* and then scaled by multiplying a constant 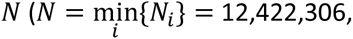 which is the minimum sample read count of all samples). The normalized read counts were then log2 transformed after adding a pseudocount 1. The normalized and log2-transformed read counts were used to represent DH levels of genomic bins.

DNase-seq data for GM12878, K562 and HSMM were downloaded from the ENCODE project (ENCODE Project Consortium 2012). The data were aligned to human genome hg19 using bowtie (Langmead et al. 2009) (http://hgdownload.cse.ucsc.edu/goldenPath/hg19/encodeDCC/wgEncodeUwDnase). The aligned reads were processed in the same way as the Epigenome Roadmap data to derive DH levels. Of note, the ENCODE data contained replicate samples for each cell type. The normalized read counts from replicate samples were first averaged to characterize the DH level for each bin in each cell type. The DH level was then log2 transformed after adding a pseudocount 1.

### Genomic loci filtering

Since most genomic loci are noise rather than regulatory elements, we filtered genomic loci to exclude those without strong DH signal in any Epigenome Roadmap DNase-seq sample. The filtering was done in three steps. First, genomic bins with normalized read count ≤8 in all samples were excluded. Second, bins with normalized read count larger than 10,000 in ≥1 sample were considered abnormal and therefore also excluded. Third, a signal-to-noise ratio (SNR) was computed for each bin in each sample, and bins with SNR ≤ 2 in all samples were considered as noise and filtered out. In order to compute SNR of a genomic bin in a sample, we first collected 500 bins in the neighborhood of the bin in question. The average DH level of these bins was computed and then log2 transformed after adding a pseudocount 1 to serve as the background. The log_2_(SNR) was defined as the difference between the normalized and log2 transformed DH level of the bin in question and the background. SNR ≤ 2 is equivalent to log_2_(SNR) ≤ 1.

After filtering, 1,136,465 genomic bins (called DNase I hypersensitive sites, or DHSs, hereinafter) with unambiguous DNase-seq signal in at least one sample were identified. All analyses in this study were performed on these genomic loci, except for the leave-one-out cross-validation analysis in **Figure 1d-i** which will be described in a separate section below.

### Bulk RNA-seq data processing

The aligned RNA-seq data (alignment based on hg19) for the same 70 Epigenome Roadmap samples were downloaded from ftp://ftp.genboree.org/EpigenomeAtlas/Current-Release/experiment-sample/mRNA-Seq/. Cufflinks (Trapnell et al. 2010) was used to compute the expression values (i.e., FPKM: fragments per kilobase of exon per million mapped fragments) using gene annotations in GENCODE (Harrow et al. 2012) (Release 19 (GRCh37.p13)). 37,335 transcripts (called “genes” hereinafter for simplicity) with FPKM > 1 in at least one sample were identified. These FPKM values were log2 transformed after adding a pseudocount 1 and then quantile normalized across samples. After normalization, the quantiles of the Epignome Roadmap training data were stored for future use. When new RNA-seq samples need to be analyzed, they will be quantile normalized against these stored quantiles.

For evaluation, we downloaded the following data from GEO: (1) GM12878 and K562 bulk RNA-seq data (GSM958728, GSM958729), (2) GM12878 RNA-seq data from small-cell-number samples with 10, 30 and 100 cells (GSM1087860, GSM1087861, GSM1087858, GSM1087859, GSM1087856, GSM1087857). For these samples, reads were mapped to human genome hg19 using Tophat (Kim et al. 2013). Gene expression values were then computed using Cufflinks in the same way as how we processed the Epigenome Roadmap RNA-seq data. Finally the gene expression values were quantile normalized with the Epigenome Roadmap RNA-seq data using the stored quantiles.

### BIRD Model

The BIRD (BIg data Regression for predicting DNase I hypersensitivity) algorithm is described and systematically evaluated in a companion article. For readers’ convenience, we review its workflow in **Supplementary Methods**. Readers are referred to Zhou *et al. (Zhou et al. submitted)* for more details. BIRD software is available at https://github.com/WeiqiangZhou/BIRD. Models trained using the 70 Epigenome Roadmap samples have been stored in the software package.

### Prediction performance evaluation

Three statistics were used in this article for evaluating prediction accuracy in different analyses. Let 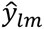 be the predicted DH level of locus *l* (=1,…, *L*) in test sample *m* (=1,…, *M*), and let *y_lm_* be the true DH level measured by DNase-seq. The three statistics include:

1. Cross-locus correlation (*r_L_*). This is the Pearson’s correlation between the predicted signals 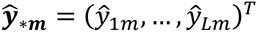 and the true signals ***y****_*m_* = (*y*_1_*_m_*,…, *y_Lm_*)*^T^* across different loci for each test sample *m*. The cross-locus correlation measures the extent to which the DH signal within each sample can be predicted.
2. Cross-sample correlation (*r_C_*). This is the Pearson’s correlation between the predicted signals 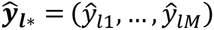 and the true signals ***y****_l*_* = (*y_l_*_1_,…, *y_lM_*) across different samples for each locus *l*. The cross-sample correlation measures how much of the DH variation across samples can be predicted.
3. Squared prediction error (*τ*). This is measured by the total squared prediction error scaled by the total DH data variance in the test dataset: 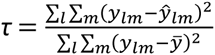, where 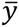 is the mean of *y_lm_* across all DHSs and test samples.

### Leave-one-out cross-validation

Leave-one-out cross-validation was used to evaluate BIRD prediction accuracy when bulk RNA-seq data were used as predictors. In each fold of the cross-validation, the 30 Epigenome Roadmap cell types (consisting of 70 samples) were partitioned into a training dataset with 29 cell types and a test dataset with 1 cell type. In other words, all samples from one cell type were used as test data, and all samples from the remaining 29 cell types were used as the training data. BIRD was then trained using all samples in the training dataset and applied to predict DH for all samples in the test dataset.

To ensure that the test data are not used in the construction of prediction models, the predictor and genomic loci filtering procedure was applied to each fold of cross-validation by using the training data only. For instance, we identified genes with FPKM > 1 in at least one RNA-seq sample in the training data as predictors. The identified predictors were a subset of the 37,335 genes described before (note: the 37,335 genes were identified using all 70 samples rather than using only the training samples). For different folds of cross-validation, a slightly different set of predictors was identified. Similarly, genomic loci (i.e. DHSs) to be predicted were selected by applying the previously described filtering protocol to the training data only: (1) normalized bin read count ≥8 in at least one sample; (2) normalized bin read count < 10,000 in all samples; (3) SNR ≥ 2 in at least one sample. Prediction models were constructed for the identified genomic loci. These loci varied from fold to fold, and they were also slightly different from the 1,136,465 genomic loci derived from all 70 samples. Of note, the parameters of BIRD (i.e. *K* and *N* in **Supplementary Methods)** were selected following the same procedure described in **Supplementary Methods** using 1% loci randomly chosen from the training dataset (i.e., samples from 29 cell types in each fold). Since the training data were different in each fold, the parameters also varied from fold to fold.

After predictions were made for all samples, *r_L_, r_C_* and *τ* were calculated between the true and predicted DH profiles. Conceptually, one can organize the predicted values into a matrix. Rows of the matrix correspond to genomic loci, and columns of the matrix correspond to samples. The matrix has missing values as not all genomic loci have prediction models in all samples. This is because genomic loci filtering was dependent on the training data. As a result, in each fold of cross-validation, prediction models were built for a slightly different set of genomic loci. To compute *r_L_, r_C_* and *τ*, missing data points in the prediction matrix were excluded, and only data points with predicted DH values were used. This produces **Figure 1d-f**.

### Random prediction models by permutation

To construct random prediction models, sample labels of the DNase-seq data in the training dataset were shuffled. This permutation broke the connection between DNase-seq and RNA-seq samples. Then, BIRD was trained by the permuted training dataset and applied to predict DH in the test dataset. The permutation was performed in each fold of the leave-one-out cross-validation and the prediction performance was then evaluated by *r_L_*, *r_C_* and *τ*. Of note, our permutation here did not perturb the locus effects of DH profile. Therefore, predictions from the random prediction models mostly captured the average DH level of each genomic locus in the training dataset.

### Wilcoxon signed-rank test

Two-sided Wilcoxon signed-rank test (Wilcoxon 1945) was performed to obtain *p*-values for comparing prediction accuracy of BIRD and random prediction models. In order to test whether two methods perform equally in terms of *r_L_*, the paired *r_L_* values from these two methods for each sample was obtained. Then the *r_L_* pairs from all samples are used for Wilcoxon signed-rank test. Similarly, to compare two methods in terms of *r_C_*, the paired *r_C_* values for each locus were obtained, and *r_C_* pairs from all genomic loci were used for the Wilcoxon signed-rank test.

### Categorization of genomic loci based on cross-sample variability

When studying the cross-sample prediction performance (i.e., *r_C_*) in **Figure 1g-h**, genomic loci were grouped into different categories based on their cross-sample variability of the predicted DH profile. First, loci with predicted DH value (at log2 scale) smaller than 2 across all cell types were treated as noisy loci (**Fig. 1g-h**, indicated by “Noisy loci”). For such loci, the observed DH level may contain substantial noise, and the cross-sample correlation between the predicted and the true DH is expected to be low (since the correlation between random noise and another independent random variable is expected to be zero). After excluding the noisy loci, the other loci were then categorized based on the coefficient of variation (CV) of the cross-sample DH values. For each locus, CV was calculated as the ratio of the standard deviation to mean of the predicted DH at this locus across all samples. Loci were divided into three categories: CV≤0.2, 0.2<CV≤0.4, CV>0.4 (**Fig. 1g-h**). A large CV indicates that the DH of a locus has more variation across samples. **Figure 1g** shows the distribution of *r_C_*. Genomic loci are grouped into bins based on *r_C_* values. For each bin, the number of loci in different CV categories is shown. **Figure 1h** shows the distribution of *r_C_* in each CV category.

### Chromatin accessibility prediction for clusters of co-activated DHSs

A transcriptional regulation process often involves co-activation of multiple *cis*-regulatory elements. Such co-activated regulatory elements can be viewed as regulatory “pathways”. Previously, DHSs discovered from the ENCODE DNase-seq data have been clustered into 2500 clusters based on their cross-sample co-variation patterns (Sheffield et al. 2013). Using these pre-defined clusters as “pathways”, we investigated how accurate the pathway activity can be predicted using bulk RNA-seq. To do so, we first identified the cluster membership of the 1,136,465 genomic loci studied here based on the clustering results provided by Sheffield et al. (2013) (obtained from http://big.databio.org/RED/TableS03-dhs-to-cluster.txt.tar.gz, the “original cluster” assignment was used). For each cluster, we then computed its mean DH level (missing values were excluded) in each sample using all DHSs in the cluster. Next, we built prediction models to predict the mean DH level of each cluster (i.e., “pathway activity”) in the same way as 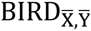. The prediction accuracy was evaluated using leave-one-out cross-validation (i.e., using 29 cell types as training and 1 cell type as test data). After cross-validation, the average DH level for each DHS cluster and each sample was obtained, and the cross-sample correlation was calculated between the true and predicted mean DH level for each cluster (**Fig. 1i**).

### Transcription factor binding site prediction

BIRD models trained using the 70 Epigenome Roadmap samples were applied to predict binding sites for 34 TFs in GM12878 cells and 25 TFs in K562 cells. For each TF, DNA motif obtained from TRANSFAC (Matys et al. 2006) and JASPAR (Mathelier et al. 2014) (**Supplementary Table 2**) was computationally mapped to the human genome using CisGenome (Ji et al. 2008) (using default likelihood ratio ≥ 500 cutoff). DHSs (i.e., the 1,136,465 genomic bins) that overlapped with motif sites were retained for subsequent analyses. These motif-containing DHSs were ranked in decreasing order based on the predicted DH level to serve as the predicted TFBSs. As a comparison, DHSs were also ranked based on two other methods: the true DH level at each DHS from the corresponding DNase-seq data (“True”) and the DH level predicted based on the mean DH profile of the 70 training samples (“Mean”).

To evaluate the prediction performance of different methods, the transcription factor ChIP-seq uniform peaks data for the 34 TFs in GM12878 and 25 TFs in K562 were downloaded from the ENCODE project to serve as the truth (http://hgdownload.cse.ucsc.edu/goldenPath/hg19/encodeDCC/wgEncodeAwgTfbsUniform/). ChIP-seq peaks overlapped with motif sites were used as the gold standard. The percentage of these gold standard peaks that were recovered by the top ranked predicted TFBSs was computed to measure the sensitivity of each prediction method. Different methods were compared by plotting the sensitivity as a function of the number of predicted TFBSs (**Fig. 1j-k, Supplementary Figs. 5-6**).

To evaluate statistical significance of the predicted TFBSs, the same BIRD models were applied to a set of randomly sampled genomic bins (n= 984,213, sampled from non-repeat genomic regions) to make predictions. Using the predicted DH values in the random genomic loci as the null distribution, a *p*-value was computed for each studied DHS to evaluate the significance of its predicted DH level (*p*-value of the predicted DH level at a DHS = [no. of random loci with equal or larger predicted DH levels] / [the total no. of random loci]). To adjust for multiple testing, the *p*-values were converted to *q*-values based on the previously described method (Dabney and Storey). *q*-values for BIRD predictions were labeled on top of each sensitivity-rank plot (e.g., **Fig. 1j-k**).

To generate **Figure 3g** and **Supplementary Figure 15** that compare different TFBS prediction methods using small-cell-number or single-cell RNA-seq data, we computed the area under the curve (AUC) for each method using the sensitivity rank curves in **Figure 3a-f, Supplementary Figure 8, Figure 5a-f**, and **Supplementary Figures 12-14**. The AUC of each method was then scaled by (i.e., divided by) the AUC obtained using the true DNase-seq data (**Supplementary Tables 3-4**). To show a clear comparison of different methods for predicting binding sites of each TF, colors in the heatmap (**Fig. 3g** and **Supplementary Fig. 15**) reflect the transformed AUC values. For instance, values within each row were transformed to the range between 0 and 1 by: [AUC value − min(value)]/[max(value) − min(value)], here min(value) and max(value) represent the minimum and maximum AUC value within each row. Of note, within each TF, the minimum AUC was transformed to 0 and the maximum AUC was transformed to 1. As a reference, the untransformed minimum AUC value from all methods was shown for each TF using a blue bar beside the TF name in **Figure 3g** and **Supplementary Figure 15**.

To measure the overall prediction performance of each method, we calculated the average rank score (shown using red bars under the name of each method in **Fig. 3g** and **Supplementary Fig. 15**) across all 34 test TFs. First, for each TF, different methods were ranked according to their AUC values. For instance, in **Figure 3g**, the best performing method has rank 1 and the worst performing method has rank 9. Then, we calculated the average rank across all test TFs for each method. Smaller average rank indicates better overall prediction performance.

### Bulk ATAC-seq data processing

ATAC-seq data for GM12878 with 50,000 and 500 cells were obtained from GEO (GSE47753). The paired-end reads were aligned to human genome hg19 using bowtie (Langmead et al. 2009) with parameters (-X2000 -m 1) which specify that paired reads (a pair of reads was referred to as a fragment) with insertion up to 2,000 base pair (bp) were allowed to align and only uniquely aligned fragments were retained. Then, PCR duplicates (i.e. fragments that aligned to exactly the same genomic location) were determined using Picard (http://broadinstitute.github.io/picard/) where only one fragment was kept and the others were removed. Next, we measured the bin-level fragment coverage by counting how many fragments covered each 200bp genomic bin. Similar to the DNase-seq data, bin-level fragment coverage for each sample was first divided by the sample’s whole-genome fragment coverage (i.e., sum of bin-level fragment coverage across the genome) and then scaled by a constant *N* (=12,422,306, to be consistent with the DNase-seq data). Finally, the normalized bin fragment coverage from different replicate samples were averaged and log2 transformed after adding a pseudocount 1.

### Histone modification ChIP-seq and MOWChIP-seq data processing

H3K27ac and H3K4me3 MOWChIP-seq data for GM12878 with 100 and 600 cells were obtained from GEO (GSE65516: GSM1666202, GSM1666203, GSM1666204, GSM1666205, GSM1666206, GSM1666207, GSM1666208, GSM1666209). For both histone marks, the processed signal files provided by the MOWChIP-seq authors were downloaded from GEO. These files contained normalized read counts for the whole genome divided by 100 bp bins (Cao et al. 2015). The data were converted to 200 bp resolution by merging adjacent two 100 bp bins (i.e., adding the read counts of the two 100 bp bins).

Due to nucleosome displacement, the spatial distribution of histone modification signal surrounding each regulatory element (e.g., transcription factor binding site) may differ from the peak of the DNase-seq and ATAC-seq signal (He et al. 2010). Therefore we first explored different ways to summarize the histone modification signal in order to maximize its correlation with the bulk DNase-seq data. To do so, we considered a W-bp long window centered at each genomic locus (200bp bin). The normalized read counts of all 200bp bins covered by the window were averaged to serve as the summary of the histone modification signal at the locus. The summarized signals from replicate samples were averaged and log2 transformed after adding pseudocount 1. We then tested different window sizes (*W*=200, 600, 1000, 1400, 1800, 2200, 2600 bp) to find the optimal *W* that maximizes the correlation between the summarized histone modification signal and the bulk DNase-seq signal (i.e., DH level at 200-bp resolution as described before) across all genomic loci (**Supplementary Fig. 9a**). For H3K27ac with 100 and 600 cells, the optimal *W* was 2200. For H3K4me3 with 100 cells, the optimal *W* was 2200. For H3K4me3 with 600 cells, the optimal *W* was 1800. The summarized MOWChIP-seq signals using these optimal *W* were then compared with BIRD and ATAC-seq in **Figure 2** and **Supplementary Figure 10**. H3K27ac and H3K4me3 ChIP-seq data for bulk GM12878 samples were obtained from ENCODE (http://hgdownload.cse.ucsc.edu/goldenPath/hg19/encodeDCC/wgEncodeBroadHistone/). For each 200bp genomic bin, reads from the bulk ChIP and input control samples were counted. Bin read counts in each sample were normalized by the sample’s total read count and then scaled by multiplying 1,000,000. Signals were then calculated as the difference between the ChIP and input samples. Similar to MOWChIP-seq, we used the average signal of a *W*-bp long window (*W*=200, 600, 1000, 1400, 1800, 2200, 2600 bp) centered at each genomic locus to represent its summarized histone modification signal. The summarized signals from replicate samples were then averaged and log2 transformed after adding pseudocount 1. The optimal *W* was determined by maximizing the correlation with the bulk DNase-seq signal. For bulk H3K27ac and H3K4me3, the optimal *W* was 1000 and 1400 respectively (**Supplementary Fig. 9c**). The summarized ChIP-seq signals using these optimal *W* were then used for generating **Supplementary Figure 10**.

For TFBS prediction using MOWChIP-seq data, we also first optimized the window size *W* for each MOWChIP-seq dataset. For each histone mark and cell number, we obtained the scaled AUC (scaling is done by dividing the AUC of using true DNase-seq data to predict TFBSs) for all 34 TFs using different window sizes. For each TF, different window sizes were then ranked based on the scaled AUC. The average rank of each window size *W* across all 34 TFs was computed (**Supplementary Fig. 9b**). *W* with the best average rank was identified. For H3K27ac MOWChIP-seq with 100 and 600 cells, the optimal *W* was 1800. For H3K4me3 MOWChIP-seq with 100 cells, the optimal *W* was 2200. For H3K4me3 MOWChIP-seq with 600 cells, the optimal *W* was 1800. The summarized MOWChIP-seq signals using these optimal *W* were then compared with BIRD and ATAC-seq in **Figure 3** and **Supplementary Figure 8**. We note that since TFBSs of different TFs may be associated with different histone modification signatures (e.g., different types of histone modifications), a better way to predict TFBS might be to use multiple types of histone modification data and develop a prediction model specific for each TF. However, that would make the TFBS prediction more difficult to apply in reality because one would need to collect more MOWChIP-seq data and have knowledge on the histone modification signature for each TF. For this reason, our analyses here were primarily focused on evaluating the performance of using one data type (i.e., H3K27ac, H3K4me3, ATAC-seq, or RNA-seq) and a common prediction procedure for all TFs. This makes the comparison among MOWChIP-seq, ATAC-seq (ATAC-b500) and BIRD (BIRD-b30) relatively fair in the sense that different TFBS prediction methods have similar level of complexity in terms of data collection and computational analysis.

### Chromatin accessibility prediction based on single-cell RNA-seq data

We downloaded two datasets from GEO: (1) GM12878 single-cell RNA-seq data (GSE44618, 28 cells in total), (2) HSMM single-cell RNA-seq data (GSE52529, 69 cells from undifferentiated HSMM were used for our analysis). For these samples, reads were mapped to human genome hg19 using Tophat (Kim et al. 2013). Gene expression values were then computed using Cufflinks in the same way as how we processed the Epigenome Roadmap RNA-seq data. For each dataset, we randomly sampled *k* cells (*k* = 1, 5, 10, 20, 28 for GM12878; *k*= 1, 5, 10, 20, 30, 40, 50, 69 for HSMM) and calculated their average gene expression profile. The average gene expression profile was then used as the input for BIRD to predict the DH profile. This is the “approach 1” in **Figure 5g-h**. For each *k* (except for *k* = 1 and 28 for GM12878, and *k* = 1 and 69 for HSMM), the random sampling was repeated 10 times. The mean and standard deviation (SD) of the results from the 10 analyses were shown in **Figures 4b** and **5g-h**. For *k*=1, the analysis was performed for every single cell.

We also tried a second approach to predict DH using single-cell RNA-seq (i.e., “approach 2” in **Fig. 5g-h**). In this approach, we first applied BIRD to predict DH for every single cell using the single-cell RNA-seq data. Then, we pooled a random group of *k* cells (note: the same cells sampled in “approach 1” were used to keep the comparison consistent) and computed the average of their predicted DH profile. The random sampling was repeated 10 times as above, and the mean and SD of the prediction performance from the 10 analyses were shown in **Figure 5g-h**. A comparison between approach 1 and approach 2 shows that using multiple cells’ average expression as predictor had much higher prediction accuracy than using each cell’s expression to make prediction and then average the predicted DH profile.

### Chromatin accessibility based on single-cell ATAC-seq data

Two single-cell ATAC-seq datasets for GM12878 were obtained. Dataset 1 (ATAC1) was obtained from GEO (GSM1647121). This dataset was a mixture of human GM12878 cells and mouse Patski cells. Paired-end reads were trimmed by Trimmomatic (Bolger et al. 2014) to remove adaptor content and aligned to human genome hg19 using bowtie2 (Langmead and Salzberg 2012) with parameter -X2000. PCR duplicates were removed using Picard. The aligned reads were then assigned to individual cells based on the barcode information and only GM12878 cells were retained for subsequent analyses. For each cell, bin-level fragment coverage was obtained for each genomic locus (i.e., 200bp bin), and bin fragment coverage was normalized in the same way as the bulk ATAC-seq data. The single-cell ATAC-seq data are highly discrete. According to the original report describing this data (Cusanovich et al. 2015), the sequencing has reached saturation and the median value of total read counts per cell was 2503. We identified GM12878 cells (n=222) with more than 500 non-zero-coverage loci and used them for the subsequent analyses. Dataset 2 (i.e. ATAC2) was obtained from GEO (GSE65360). This dataset contains GM12878 ATAC-seq for 384 single cells. For each single cell, paired-end reads were trimmed by Trimmomatic to remove adaptor content and aligned to human genome hg19 using bowtie2 with parameter -X2000. PCR duplicates were removed using Picard. Then, bin-level fragment coverage for each cell was computed, normalized and transformed in the same way as single-cell ATAC-seq dataset 1. 340 cells with more than 500 non-zero-coverage loci were retained for the subsequent analyses.

For the single-cell ATAC-seq dataset 1, we randomly sampled a group of *k* cells (*k* = 1, 5, 10, 20, 28, 50, 100, 222) and calculated their average ATAC-seq profile (i.e., average of the normalized bin fragment coverage). The average profile was then log2-transformed after adding pseudocount 1. For each *k* (except for *k*=1 and 222), we repeated the random sampling 10 times. The mean and SD of results from the 10 analyses were shown in **Figure 4b**. For *k*=1, the analysis was performed on every single cell. The same analysis was also performed for the single-cell ATAC-seq dataset 2 with *k* cells (*k* = 1, 5, 10, 20, 28, 50, 100, 222 and 340).

### Hybrid prediction based on combining single-cell RNA-seq and single-cell ATAC-seq

For the hybrid approach, we randomly sampled *x* (*x* = 22, 72, 194 and 312) cells from the single-cell ATAC-seq dataset 2. Dataset 2 was used since it performed better than dataset 1 based on analyses in **Figure 4b**. We obtained the average ATAC-seq profile of the sampled cells using the protocol described above. We also obtained BIRD-predicted DH from pooled single-cell RNA-seq using 28 cells. The average of the ATAC-seq profile and BIRD predicted DH profile was then computed. The total number of cells used by this hybrid approach was *k* = *x*+28 (i.e., *k* = 50, 100, 222 and 340). In **Figure 4b**, this hybrid approach was compared to pooled single-cell ATAC-seq using the same number of cells. For the hybrid approach, the sampling of cells from scATAC-seq was repeated 10 times. The mean and SD of results from the 10 analyses were shown in **Figure 4b**.

## ACKNOWLEDGMENTS

This research is supported by grants from the National Institutes of Health (R01HG006282 and R01HG006841) and a Seed Fund of the Johns Hopkins Institute for Data Intensive Engineering and Science.

